# Obesity Impairs the Antitumor Activity of CAR-T Cells in Triple-Negative Breast Cancer

**DOI:** 10.1101/2025.10.10.681694

**Authors:** Hannah M. Malian, Caroline Marnata Pellegry, Hannah M. Oh, Elaine M. Glenny, Alyssa N. Ho, Gianpietro Dotti, Stephen D. Hursting, Michael F. Coleman

## Abstract

**Background:** We have reported that chimeric antigen receptor (CAR) T cells targeting B7-H3 (B7-H3.CAR) are effective in a preclinical model of triple negative breast cancer (TNBC), and have initiated a Phase I study to assess safety and efficacy. However, heterogeneous antigen expression and immunosuppressive tumor microenvironments (TME) remain roadblocks for effective CAR-T cell therapy. In particular, obesity represents a negative prognostic factor in TNBC partly due to chronic inflammation and impaired adaptive immune responses. Hence, we sought to determine if obesity can affect the antitumor activity of B7-H3.CAR-T cells.

**Methods:** We used qPCR and western blotting to determine if cytokines associated with obesity affect B7-H3 expression in TNBC cell lines. Furthermore, we used shRNA to suppress B7-H3 expression in a syngeneic orthotopic E0771 tumor model and measured tumor growth in control and diet-induced obese (DIO) mice. Finally, we evaluated the antitumor effects of B7-H3.CAR-T cells in both control and DIO mice orthotopically engrafted with the E0771 tumor cell line. Immune profiling was conducted using flow cytometry.

**Results:** Obesity-related inflammatory cytokines promote B7-H3 expression in human and murine TNBC cells in vitro and B7-H3 expression correlates with tumor aggressiveness in vivo. CAR-T cells obtained from control or DIO mice were equally cytotoxic in vitro but activated T cells and B7-H3.CAR-T cells obtained from DIO mice show transcriptomic changes (enriched *Tox2, Prdm1, Batf*) and impaired glycolytic capacity, respectively. Finally, we demonstrated that obesity impairs CAR-T cell antitumor effects and durability of response in vivo with a near complete loss of memory formation.

**Conclusions:** Here we identified a correlation between B7-H3 expression, obesity, and rate of tumor growth in TNBC. Furthermore, we showed that obesity constrains both the ability of B7-H3.CAR-T cells to control tumor growth and to elicit durable immunological memory. Taken together, these data identify obesity as an underappreciated and potent modulator of CAR-T cell functionality.

**What is already known:** B7-H3 protein is upregulated in many human malignancies including TNBC and is often associated with worsened outcomes. B7-H3.CAR-T cells show promise in preclinical models of TNBC and entered clinical translation.

**What this study adds:** This study identifies a previously unknown correlation between obesity, B7-H3 expression, and rate of tumor growth in TNBC. Furthermore, our preclinical model of B7-H3.CAR-T cell therapy demonstrates that obesity negatively affects the antitumor activity of B7-H3.CAR-T cells in TNBC.

**How this study affects other research/practice:** This study highlights obesity as an understudied and critically important covariate for adoptive T-cell therapy and demonstrates important links between systemic metabolism and antigen expression. This work paves the way for future mechanistic and translational research into how obesity impacts CAR-T cell functionality.

## Introduction

Triple-negative breast cancer (TNBC) is notoriously difficult to treat and has a meager 5-year survival rate of 12% once metastasized.(1–4) TNBC’s aggressiveness stems partly from its ability to evade immune surveillance through immunosuppressive mechanisms,(5) and targeting immune checkpoints with checkpoint blockades (e.g. pembrolizumab for PDL1) has significantly improved outcomes.(6)

Adoptive transfer of CAR-T cells is an emerging therapeutic strategy for solid tumors including TNBC. Specifically for TNBC, we and others have identified B7-H3 (CD276) protein as a potential target for CAR-T cells.(7,8) B7-H3 is expressed in 50-80% of TNBC tumors,(8–10) and high B7-H3 expression is associated with worsened survival.(11) B7-H3 has limited expression in normal adult tissue,(7) which makes it a promising target for CAR-T cell therapy. Thus, we have developed B7-H3.CAR-T cells and demonstrated activity and safety in multiple preclinical solid tumor models, including TNBC.(7,8,12–14) Based on these findings, early phase clinical trials to assess safety and efficacy of B7-H3.CAR-T cells have been initiated in solid tumors including in patients with TNBC.(15)

Obesity is associated with higher incidence of TNBC and promotes faster tumor progression.(16,17) While the mechanistic links between obesity and cancer remain incompletely understood, immunosuppressive tumor micro environments (TME)(18–20) and T cell dysfunction(18,19,21–23) are thought to be two driving factors. However, obesity seems to predict a paradoxically favorable response to immune checkpoint inhibitor treatment across many cancers.(24–26) Therefore, given the high and rising global rates of obesity,(27) understanding if obesity-associated immune impairments can impact the therapeutic efficacy and persistence of CAR-T cells is critical.

In this study, we developed in vitro and in vivo TNBC models to study the effects of obesity-associated cytokines and diet-induced obesity on the expression of the target antigen B7-H3 and adoptive transfer of B7-H3.CAR-T cells.

## Methods

### TNBC cell culture models

MDA-MB-468, MDA-MB-231, and E0771 cells were cultured in Dulbecco’s Modified Eagle’s Medium (DMEM) (Gibco) with 10% fetal bovine serum (FBS) (Corning), 1% penicillin-streptomycin (Gibco), and 2 mM L-glutamine (Gibco). B7-H3 (CD276) expression was suppressed in E0771 cells by lentiviral transduction, using particles generated by transfection of HEK293T cells with psPAX2, pMD2.G, and B7-H3-targeting shRNAs in a psi-LVRU6H vector (Genecopoeia). The target shRNA sequences were CCAATGGACTTAATTCCCATC (shB7-H3), and GCTTCGCGCCGTAGTCTTA (scramble). E0771 cells were selected with hygromycin (500 µg/ml) to enrich for successfully transfected cells. B7-H3 (CD276) was overexpressed in E0771 cells via transduction with a lentiviral vector (pCDH) encoding human B7-H3. Cells (E0771_B7-H3HI) were sorted for high expression of B7-H3 using a BD FACSAria III at the UNC Flow Core Facility. All cell lines were tested for mycoplasma using the Universal Mycoplasma Detection Kit (ATCC #30-1012K).

### Cytokine treatments

Tumor cells were treated with the cytokines TNF or IFNγ, or a combination of TNF and IFNγ at 20 ng/mL for 24 h in DMEM with 1% penicillin-streptomycin, and 2 mM L-glutamine. After 24 h of cytokine treatment, cells were frozen at –80 °C until processing (except if they were used for flow cytometry, in which case cells were stained with anti-CD276 (Abcam #134161) and BV421 goat anti-rabbit IgG (BD Biosciences #565014)).

### RNA extraction, cDNA synthesis, and qPCR

RNA was isolated using the omega Biotek EZNA RNA isolation kit according to manufacturer’s instructions. Sample concentration was determined by NanoDrop UV absorbance. cDNA was synthesized with 2 µg RNA and the High-Capacity cDNA Reverse Transcription Kit according to manufacturer’s instructions (ThermoFisher Scientific, #4374967). qPCR was performed using Universal SYBR Green Supermix (Biorad # 1725271) and analyzed on a Viia 7 (ThermoFisher Scientific). Primer sequences for mouse *Cd276* (F: ACCATCACACCCCAGAGAAG R: GCCAGATGAGGTTGAGCTGT) and mouse *Rpl4* (F: CCGTCCCCTCATATCGGTGTA, R: GCATAGGGCTGTCTGTTGTTTTT) were obtained from PrimerBank and purchased from IDT (Coralville, IA). Human *CD276* and *RPL4* Taqman probes were obtained from ThermoFisher (Cat #4331182 and #4331182, respectively). RPL4 was used as a housekeeping gene for normalization. The relative difference in gene expression was calculated using the ΔΔCt method.

### Western Blot

Proteins were isolated in RIPA buffer supplemented with a protease inhibitor cocktail (Sigma Aldrich) and phosphatase inhibitors (sodium orthovanadate, sodium pyrophosphate, and β-glycerophosphate). Forty µg of protein lysate was loaded and resolved on 4-16% gradient polyacrylamide gels and transferred to nitrocellulose membranes (Bio-Rad). After blocking for 1 h in 5% BSA, membranes were incubated overnight at 4 °C with anti-CD276 antibody (1:1000, #ab134161, Abcam), anti-β-Actin, (1:5000, #sc-47778, Santa Cruz Biotechnology), or anti-phospho-P65 (1:5000, #30335, Cell Signaling). Blots were imaged using a LICOR Odyssey and analyzed with ImageStudio software (LICOR Biotechnology).

### Retrovirus preparation

Retroviral supernatant used for the transduction of activated murine T cells was generated by co-transfecting HEK293T cells with a retroviral vector containing the murine version of the B7-H3.CAR (mB7-H3.CAR) described previously,(7) and the pCL-Eco plasmid that encodes gag/pol/env using the GeneJuice transfection agent (Sigma-Aldrich). Supernatant containing the retrovirus was collected 48 and 72 hours after transfection and filtered through 0.45 μm filters.

### T cell isolation and CAR-T production

Spleens were collected from C57BL6 female mice and T cells were isolated with the MojoSort mouse CD3 T cell isolation kit (Biolegend #480024) by negative selection according to manufacturer’s instructions. T cells were cultured in Hyclone RPMI-1640 (Cytiva), with 10% Hyclone FBS (Cytiva), 1% Pen/Strep (Gibco) and 1% L Glutamax (Gibco) and 100 μM β-mercaptoethanol (Gibco, #145044). Murine T cells were stimulated for 48 hours on plates coated with 1 μg/ml anti-CD3 and 1 μg/ml CD28 (eBioscience #14-0032-82, #16-0289-81). Activated murine T cells were then transduced with retroviral supernatants using retronectin plates (Takara) and cultured for 72 h as previously described.(7) After removal from retronectin plates, murine T cells were expanded in complete Hyclone RPMI medium described above and transduction efficiency was assessed by flow cytometry using recombinant human B7-H3-Fc chimeric protein (R&D Cat# 1027-B3; 1:400) and goat-anti human IgG (H+L) antibody, Alexa Fluor® 647 (Jackson ImmunoResearch Laboratories, Inc. Cat# 109-606-088; 1:400)

### RNA-sequencing

RNA was isolated from T cells and tumors using the RNeasy micro kit (Qiagen). RNA was quantified on a Qubit 2.0 fluorometer (Life Technologies, Carlsbad, CA) and integrity was assessed on a TapeStation 4200 (Agilent Technologies, Santa Clara, CA). RNA sequencing libraries were prepared using a NEBNext Ultra II RNA library prep kit (New England Biolabs, Ipswitch, MA) with a ERCC RNA spike-in (Thermo Fisher Scientific). mRNAs were enriched and used for cDNA synthesis. Sequencing was performed on an Illumina HiSeq instrument using a 2 x 150 bp paired-end configuration at a depth of >25 million reads per sample.

### GSEA

GSEA (v4.3.3) was performed on DESeq2 normalized RNA sequencing data using the Molecular Signatures Database (MSigDB) Hallmark gene sets (v2024.1). Default settings were used to generate a normalized enrichment score for all gene sets tested.

### Flow Cytometry

Tumors harvested from tumor-bearing mice were dissociated into single cells using the Miltenyi Biotec tumor dissociation kit for mouse (#130-096-730) according to manufacturer’s protocol with enzyme R reduced to 20%. Immune cells were isolated using a Percoll gradient. Blood was collected via submandibular bleed from tumor-bearing mice into 2 mM EDTA. Blood samples were then incubated with ACK Lysing Buffer (#A1049201) before being resuspended in PBS with 2% FBS prior to staining. For immune cell isolation from mammary fat pads, single cells were isolated using the Miltenyi Biotec adipose tissue dissociation kit (#130-105-808) according to manufacturer’s protocol. All immune cells were passed through a 70 μm filter to ensure a single cell suspension prior to staining. Cells were incubated with TruStain FcX (#101319), live/dead blue viability dye, and then with an antibody cocktail of extracellular markers in BD Biosciences horizon brilliant stain buffer (#563794) and fixed with BD Biosciences cytofix/cytoperm kit (#554714). For intracellular staining, cells were permeabilized using eBioscience Foxp3/transcription factor staining buffer set (#00-5523-00) and then incubated with an antibody cocktail of intracellular markers (Supplementary Table 1). CD276 expression in TNBC cells was assessed using the anti-CD276 (Abcam #134161) and BV421 goat anti-rabbit IgG (BD #565014). All samples were analyzed using full spectrum flow cytometry with a Cytek Aurora (Cytek Biosciences) (UNC Flow Cytometry Core Facility) and analyzed using FlowJo software.

### CAR-T cell coculture

E0771_B7-H3HI tumor cells (3.0×10^4^) were cocultured with B7-H3 CAR-T cells or control non-transduced (NT) cells at a ratio of 1:1 or 1:2 in 24-well plates in the absence of cytokines. After 72 h, all cells were harvested and stained to detect either tumor cells or T cells and analyzed by flow cytometry. Counting beads (Invitrogen #C36950) were used to assess absolute numbers of cells.

### ELISA assays

In coculture experiments, supernatant was harvested after 24 h. Murine IFNγ (R&D Systems,# DY485) and TNFα (R&D Systems, cat# DY410) were quantified using commercially available ELISA kits and were conducted according to the manufacturer’s protocol.

### Seahorse assays

Seahorse Metabolic Flux Analyzer XFe96 or XFe24 instruments (Agilent Seahorse Technologies, Santa Clara, CA) were used to determine glycolytic proton efflux rate (glycoPER). B7-H3.CAR-T and control NT cells were incubated for 18 h with recombinant human B7-H3-Fc chimeric protein (R&D #1027-B3) in T cell media (Hyclone RPMI-1640 (Cytiva), with 10% Hyclone FBS (Cytiva), 1% Pen/Strep (Gibco), 1% L Glutamax (Gibco) and 100 μM β-mercaptoethanol (Gibco, #145044)). 1 h prior to analysis, 2.5×10^5^ T cells/well were seeded in triplicate and incubated at 37°C in assay media (serum-free RPMI-1640 media with 10 mM glucose, 2 mM glutamine, and 1 mM pyruvate, without bicarbonate, pH 7.4) at atmospheric CO_2_. Glycolytic rate assays were performed according to manufacturer’s protocol (Agilent).

### Mouse husbandry

Female 8-12-week-old C57BL6/NCrl mice (Charles River, Wilmington, MA) were acclimated for one week, then randomized to either a 60% high-fat diet (Research Diets D12492, New Brunswick, NJ) or an isonitrogenous matched control diet with 10% fat (Research Diets D12450J) for 15-20 weeks to generate diet-induced obese (DIO) and control (CON) phenotypes, respectively. Mice were provided ad libitum access to food and water, and were monitored by vivarium staff daily for signs of dehydration, pain, or distress. All animal studies were conducted with the approval of UNC IACUC (protocol 22-152).

### shB7-H3 in vivo experiment

Control and shB7-H3 E0771 cells (5×10^4^ cells/50 µl PBS) were orthotopically transplanted into the fourth mammary fat pad (MFP) of C57BL6/NCrl mice. Body weight was measured weekly and tumor volume (0.5 × length × width^2^) was measured for 21 days.

### B7-H3.CAR-T in vivo experiments

Mice bearing E0771_B7-H3HI tumors were randomized to CAR-T cell therapy groups after 14 days of growth. All mice then underwent lymphodepleting chemotherapy (300 mg/kg of cyclophosphamide for 1 day, 8 mg/kg fludarabine for 3 days). Forty eight hours after chemotherapy, mice received either 2.5×10^6^ control non-transduced (NT) T cells or B7-H3.CAR-T cells via tail vein injection. Survivors (defined as mice that cleared their tumors and remained tumor-free for 100 days after initial CAR-T cell treatment) were rechallenged with E0771_B7-H3HI tumor cells (5×10^4^ cells) in the 4^th^ MFP and monitored again until maximum tumor volume was reached (≥2 cm^3^ or ≥ 20 mm in any direction) or for 80 days of survival.

### Statistical Analyses

Statistical analyses were performed using GraphPad Prism (version 10.6.0) and statistically significant differences were defined as p <0.05. Differences between two groups were analyzed by Student’s t-tests or paired t-tests. Differences between three or more groups were analyzed via one-way or two-way ANOVAs (when a parametric test was appropriate) followed by Tukey’s multiple comparisons adjustment. Measurements over time were assessed using linear regression with false discovery rate (FDR) q-value correction. For differential gene expression, FDRq <0.05 was considered significant. Statistical tests and sample size are indicated in corresponding figure legends.

## Results

### Obesity-associated cytokines promote B7-H3 expression in TNBC

High expression of B7-H3 is associated with lower survival in multiple types of breast cancer, including TNBC (**Fig. S1A-C**), and obesity-driven inflammation promotes expression of immunosuppressive molecules such as PD-L1.(28,29) Hence, we sought to determine if obesity-derived inflammatory cues— such as IFNγ and TNF—might regulate B7-H3 expression. We found that IFNγ and/or TNF promoted *B7-H3* gene expression in murine (E0771 (**Fig. 1A, Supp.** Fig. 1D, E), and human (MDA-MB-231 and MDA-MB-468; **Fig. 1B-C, Supp.** Fig. 1F-I) TNBC cell lines. Similarly, TNF promoted the phosphorylation of p65 in MDA-MB-468 cells and upregulation of the expression of B7-H3 protein (**Fig. 1D-G**). Overall, these data suggest that cytokines that are increased in obesity may upregulate the expression of B7-H3 in tumor cells, potentially increasing their recognition by B7-H3.CAR-T cells.

**Figure 1.**
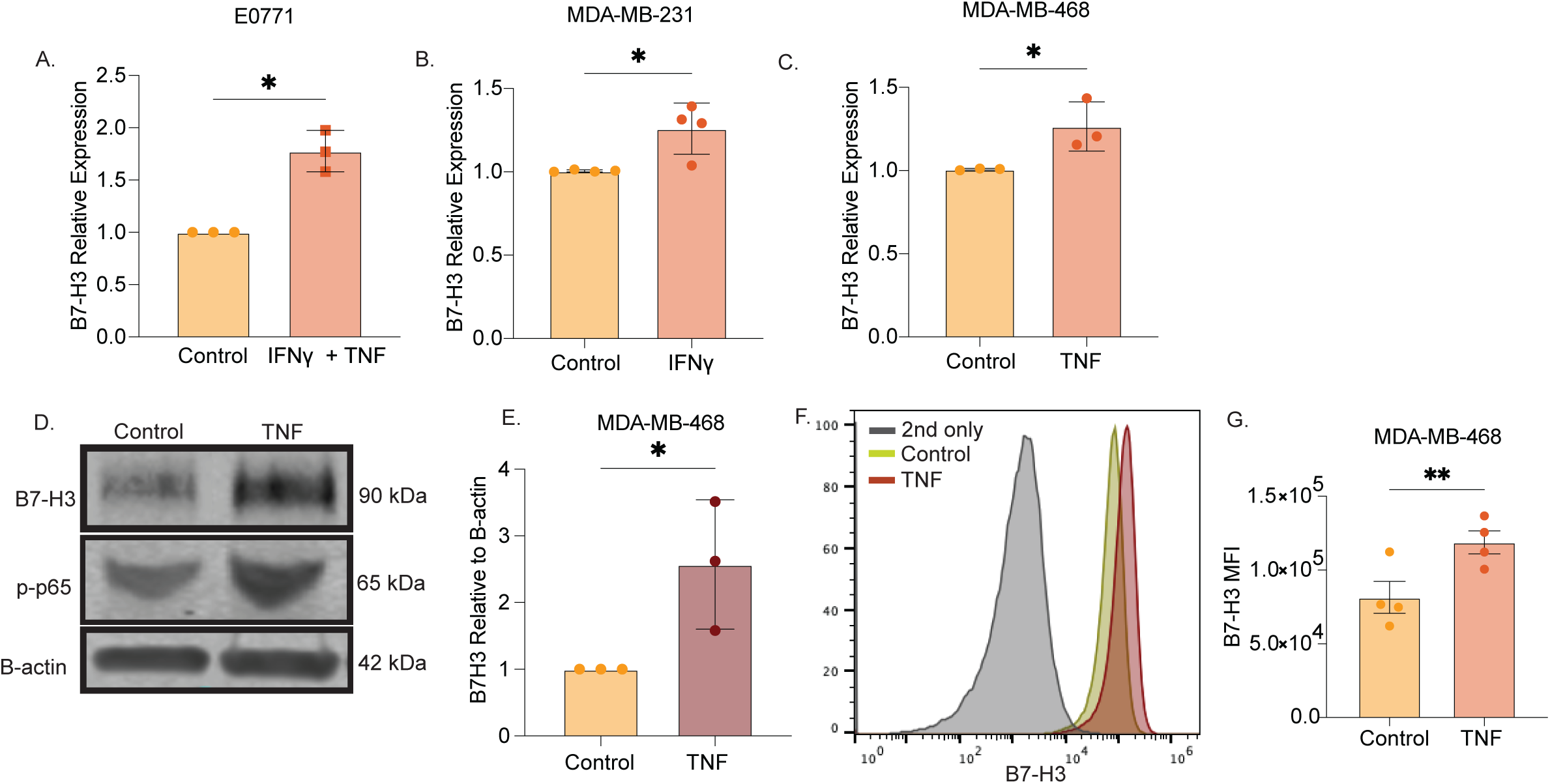
Obesity-associated cytokines promote B7-H3 expression in TNBC. A-C. qPCR quantification of *B7-H3* mRNA in one murine tumor cell line (E0771) (A) and two human tumor cell lines (MDA-MB-231 and MDA-MB-468) (B, C) treated with either TNF or TNF + IFNγ for 24 hours. D, E. Western blot quantification of the B7-H3 protein in MDA-MB-468 cells treated with TNF or control for 24 hours. F, G. Flow cytometry histograms (F) and flow cytometry mean fluorescence intensity (MFI) (G) showing surface expression of B7-H3 in MDA-MB-468 cells treated with TNF or control for 24 hours. n=3-4 group. Significance determined by Student’s t-test. *p < 0.05; **p < 0.01.

### B7-H3 expression accelerates TNBC growth in obese mice

Given our observation that obesity-associated cytokines promote B7-H3 expression, we next queried the functional contribution of B7-H3 expression in TNBC growth in obesity. Non-obese (CON) and diet-induced obese (DIO) mice (**Fig. 2A, B)** were randomized to undergo orthotopic transplantation of either B7-H3 suppressed (shB7-H3) or control (Scramble) E0771 tumor cells (**Supp.** Fig. 2**. A,B**). Absent the protumor effects of obesity, suppression of B7-H3 did not alter TNBC tumor growth (**Fig. 2C**). In contrast, suppression of B7-H3 in E0771 tumor cells significantly curtailed tumor growth in the DIO mice relative to scramble controls (**Fig. 2D**). We observed that tumor mass, T cell infiltrate, dendritic cell (DC), macrophage, and B cell numbers were similar in endpoint tumors in CON mice engrafted with either shB7-H3 or scramble E0771 tumor cells (**Fig. 2E-H, Supp.** Fig. 2C-D). In contrast, in DIO mice B7-H3 suppression resulted in significantly smaller tumors, increased intratumoral T cells, DCs, and less B cell infiltration, yet no difference in macrophages compared to scramble E0771 tumor cells (**Fig. 2I-L, Supp.** Fig. 2E,F). Finally, we performed RNA sequencing and GSEA analysis to interrogate pathways and processes altered by obesity and B7-H3 suppression. As expected, obesity sharply suppressed numerous markers of antitumor immunity. Specifically, suppression of B7-H3 expression promoted signatures of IL6/Jak/Stat3 signaling, inflammatory response, and interferon signaling in tumors from DIO mice (**Fig. 2M)**. Overall, these data indicate that suppressing B7-H3 expression in TNBC in obese mice counteracts obesity-driven immune dysfunction.

**Figure 2.**
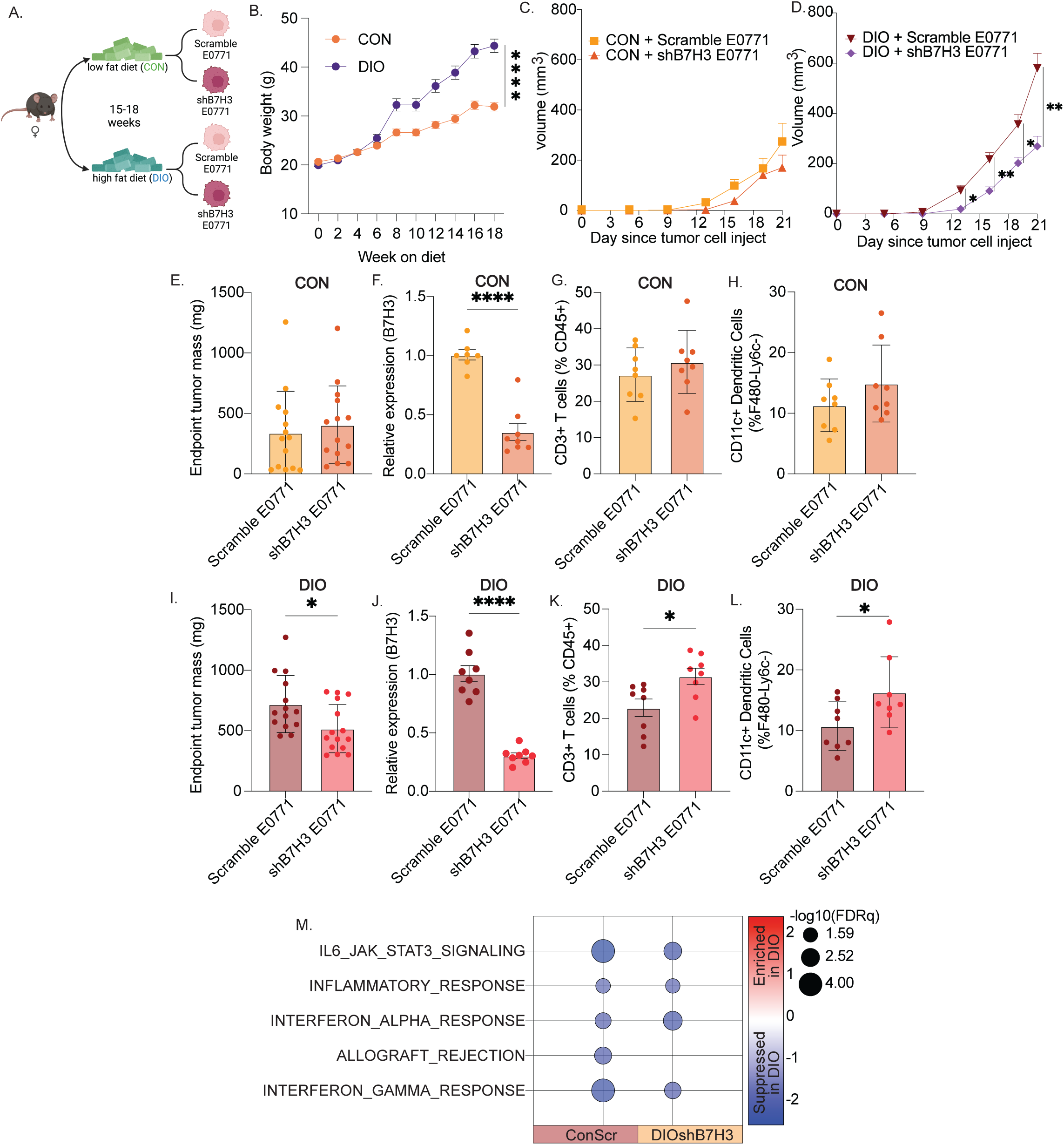
B7-H3 accelerates TNBC growth in obese mice. A. Schematic representation of the in vivo tumor model in which mice were placed on low (CON) or high fat (DIO) diet for 15-18 weeks prior to engraftment of E0771 tumor cells engineered to express either scramble shRNA (Scramble E0771) or shRNA targeting *B7-H3* (shB7H3 E0771). B. Body weight measurement over time of mice consuming low-fat diet (CON) or high-fat diet (DIO). C, D. Tumor volume measured over time in CON mice (C) and DIO mice (D). E, F. Weights (E) and *B7-H3* mRNA expression (F) in tumor harvested at the time of euthanasia in CON mice. G, H. Quantification of CD45+CD3+ T cells (G) and CD45+ CD11b+ F4/80-Ly6c-CD11c+ dendritic cells (H) in tumors collected at the time of euthanasia in CON mice as assessed by flow cytometry. I, J. Weights (I) and *B7-H3* mRNA expression (J) in tumor harvested at the time of euthanasia in DIO mice. K, L. Quantification of CD45+CD3+ T cells (K) and CD45+ CD11b+ F4/80-Ly6c-CD11c+ dendritic cells (L) in tumors collected at the time of euthanasia in DIO mice as assessed by flow cytometry. n=14/15 per group (C-E,I), n=7/8 per group (F-H, J-L). M. GSEA plot of relative hallmark gene sets enriched or suppressed in scramble E0771 tumors in DIO mice relative to scramble E0771 tumors in CON mice or shB7H3 E0771 tumors in DIO mice, n=8 mice/group. Significance determined by one-way ANOVA (B), linear regression (C,D) and Student’s t-test (E-L). *p < 0.05; **p < 0.01; ****p < 0.0001.

### B7-H3.CAR-T cells control tumor growth in a syngeneic mouse model of TNBC

B7-H3.CAR-T cells were generated from CON mice and these cells effectively targeted E0771_B7-H3HI tumor cells in in vitro coculture experiments (**Fig. 3A,B** and **Supp.** Fig. 3A,B). Cytotoxic activity of B7-H3.CAR-T cells was confirmed via release of Th1 cytokines (IFNγ and IL-2) in the culture supernatant (**Figs. 3C,D**). To evaluate the antitumor effects of B7-H3.CAR-T cells in vivo, we orthotopically transplanted E0771_ B7-H3HI into 12-week-old female C57BL/6 mice and allowed tumors to grow for 14 days before mice underwent lymphodepleting chemotherapy followed by intravenous infusion of B7-H3.CAR-T cells or control non-transduced T cells (NT). B7-H3 CAR-T cells effectively controlled tumor growth compared to mice receiving NT cells (**Fig. 3E**). Overall, these data indicate that as observed in immunodeficient models,(8) B7-H3.CAR-T cells can control tumor growth in a syngeneic tumor model of TNBC.

**Figure 3.**
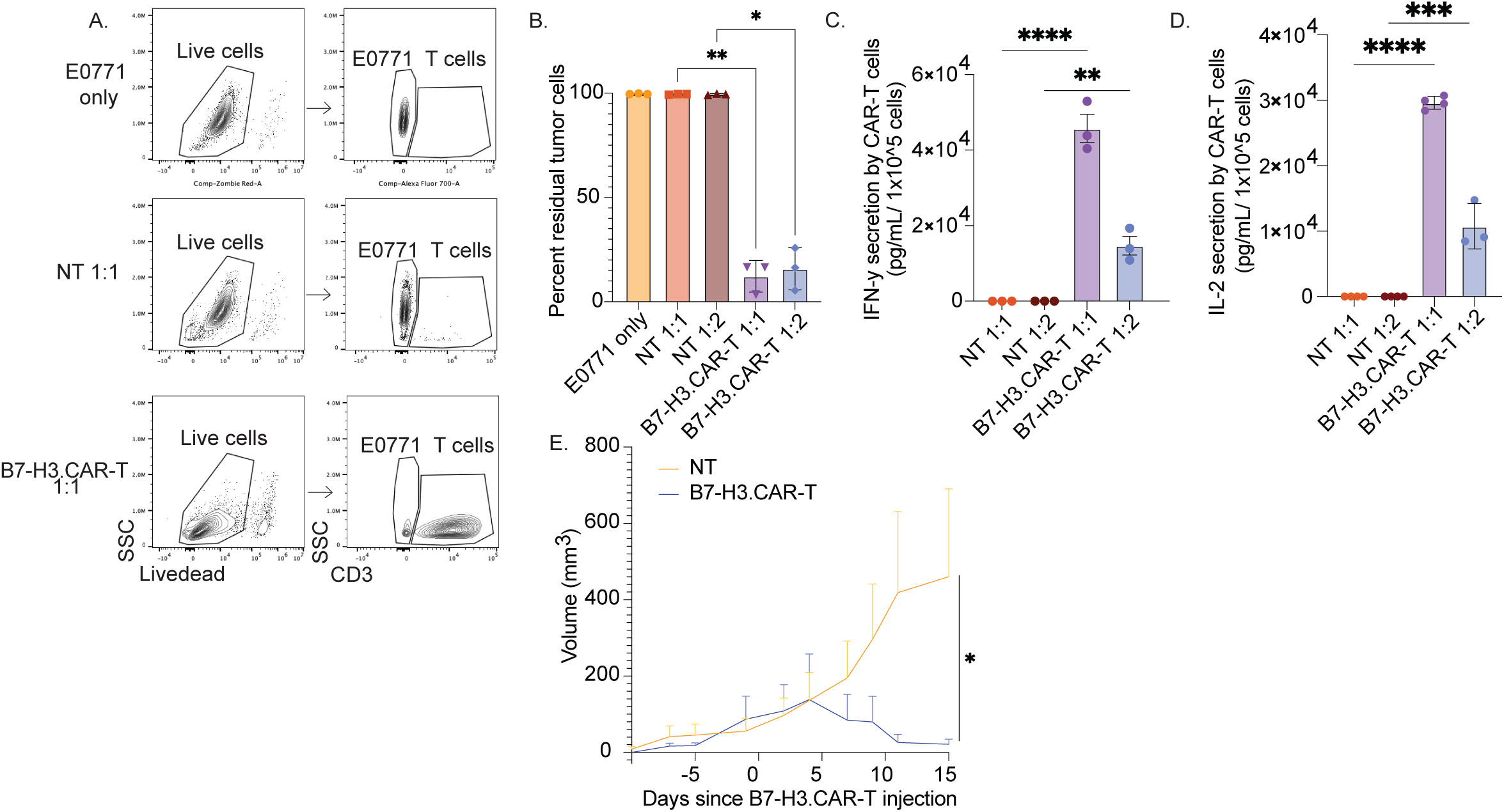
B7-H3.CAR-T cells control tumor growth in a syngeneic mouse model of TNBC. Representative flow cytometry plots (A) and summary (B) of experiments in which control (NT) and B7-H3.CAR-T cells were cocultured with the E0771 tumor cells lines. All cells were collected after 3 days of culture to quantify CD3+ T cells and tumor cells. C, D. Quantification of IFNγ (C) and IL-2 (D) by ELISA in the culture supernatant collected after 24 h in the experiments described in (A). E. Measurement of tumor volume over time in 12-week-old C57BL/6 female mice engrafted with the E0771_B7H3HI tumor cell line and treated with age-matched non-transduced T cells (NT) or B7-H3.CAR-T cells. Mice received a chemotherapy-based lymphodepletion regimen before infusion of T cells. Day 0 indicates the time of T-cell treatment. n= 3-4/group (B-D) n= 5-6 mice/group (E). Significance determined by one way ANOVA with Tukey’s multiple comparisons (B-D), and linear regression (E). *p < 0.05; **p < 0.01; ***p < 0.001; ****p < 0.0001

### CAR-T cells from DIO mice show equal effector function, but impaired metabolic fitness

We compared the functional activity of B7-H3.CAR-T cells generated from CON mice and those generated from DIO mice taking into account that obesity is known to result in functional defects in T cell cytolytic properties.(18,19,21–23) T cells isolated and activated from CON or DIO mice showed similar transduction efficiency (**Fig. 4A**), similar cytolytic activity against tumor cells (**Fig. 4B**), cytokine secretion (IFNγ, IL-2) (**Figs. 4C,D**), and granzyme-B and BCL-2 expression (**Figs 4E,F**). We also observed no significant differences in the frequency of PD1+TIM3+ cells after 3 days of coculture with tumor cells (**Fig. 4G**). However, among PD1+TIM3+ cells, we observed significantly higher frequency of LAG3+ (**Fig. 4H**) and TOX+ cells in CAR-T cells generated from DIO mice (**Fig. 4I**). Additionally, CAR-T cells from DIO mice exhibited lower CD44+CD62L+ memory cells after coculture with tumor cells compared to CAR-T from CON mice, while memory cell frequency was similar in CAR-T cells from both CON and DIO mice before antigen stimulation (**Fig. 4J,K**). Given that obesity-associated metabolic defects are well reported in T cells,(18,19,21–23) we assessed metabolic function of B7-H3.CAR-T cells using glycolytic rate assays. CAR-T cells from CON and DIO mice stimulated with recombinant B7-H3 protein showed similar basal glycolysis (**Fig. 4L, M**). Yet, when comparing compensatory glycolysis, or the ability to meet glycolytic energy demands under stress, CAR-T cells from DIO mice had significantly impaired glycolytic cell capacity (**Fig. 4N**). Overall, our data indicate that obesity does not affect the cytolytic capacity of ex vivo generated B7-H3.CAR-T cells. However, CAR-T cells generated from DIO mice might be imprinted to more rapidly become exhausted and be less effective at meeting energy demands.

**Figure 4.**
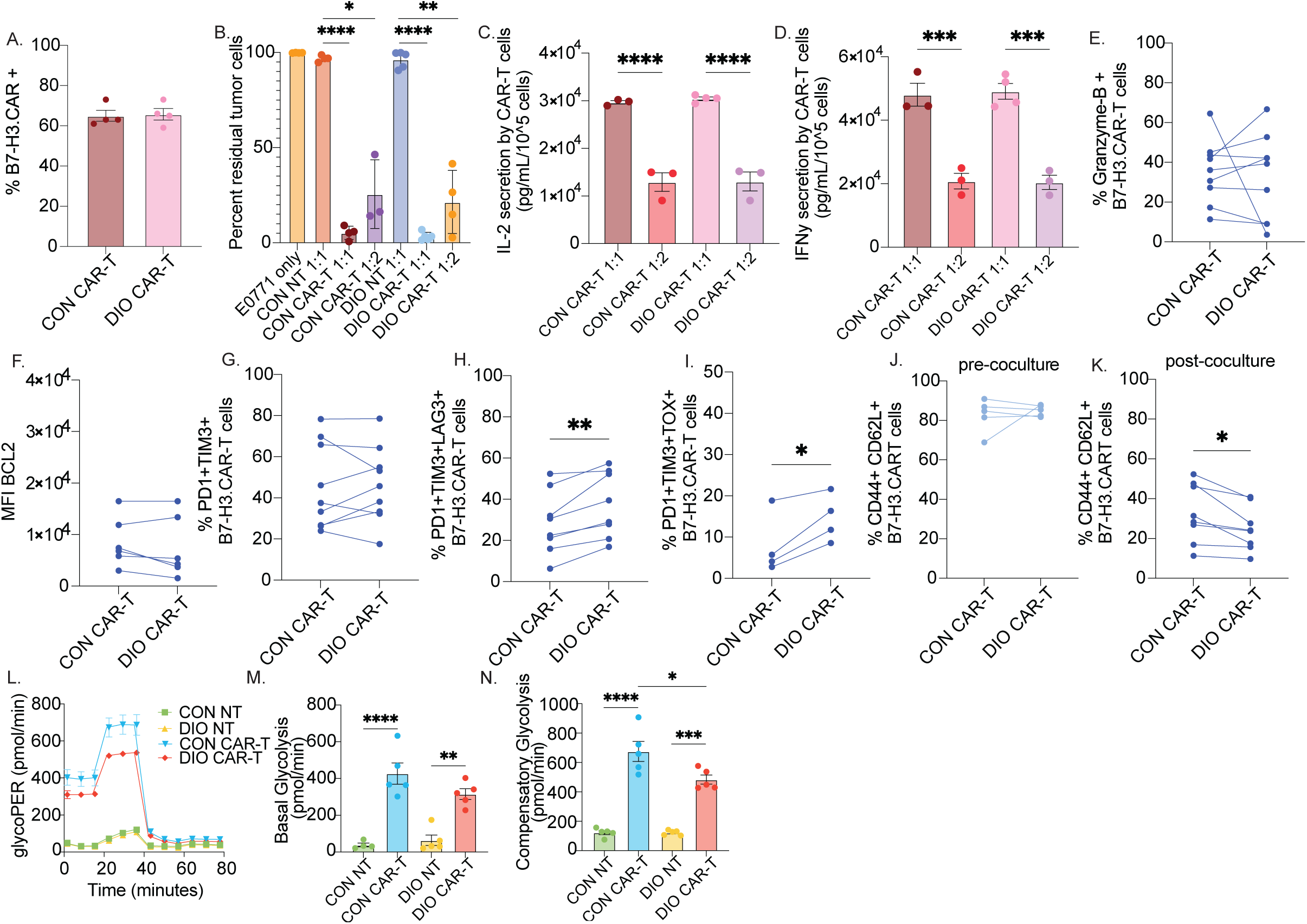
CAR-T cells from DIO mice show equal effector function, but impaired metabolic fitness. A. CAR expression in T cells generated from either CON or DIO mice as measured by flow cytometry. B. Quantification of residual tumor cells in coculture experiments in which non-transduced T cells (NT) or B7-H3.CAR-T cells generated from either CON or DIO mice were cocultured with the E0771_B7H3HI tumor cell line for 3 days. C, D. Quantification of IL-2 (C) and IFNγ (D) by ELISA in the supernatants collected after 24 h in the experiments described in (B). E-I. Immune profiling of B7-H3.CAR-T cells obtained from CON or DIO mice was performed after 3 days of coculture with tumor cells to quantify the frequency of cells expressing Granzyme-B (E), BCL2^High^ (F) and B7-H3.CAR-T cells co-expressing PD1 and TIM3 (G), PD1, TIM3 and LAG3 (H), and PD1, TIM3 and TOX (I). J, K. Co-expression of CD44 and CD62L in B7-H3.CAR-T cells before (J) and 3 days after coculture (K) with the E0771_B7H3HI tumor cell line. L. Representative glycolytic rate assay of non-transduced T cells (NT) or B7-H3.CAR-T cells, generated from either CON or DIO mice and stimulated for 18 h with recombinant B7-H3-Fc chimeric protein. M, N. Quantification of basal glycolysis (M) and compensatory glycolysis (N). n=4-6/group (A-D, F, I-J, L-N) n=8-9/group (E,G-H,K). Significance determined by Student’s t-tests (A, E-K), one-way ANOVA with Tukey’s multiple comparisons (B-D), and two-way ANOVA with Tukey’s multiple comparisons (M, N). *p < 0.05; **p < 0.01; ***p < 0.001; ****p < 0.0001.

### Activated T cells from DIO mice reveal a premature effector signature

We sought to understand how obesity may dysregulate T cell differentiation toward exhaustion and memory states, and metabolic fitness, by exploring the transcriptome of recently activated T cells. T cells were isolated from spleens of CON and DIO mice and activated with anti-CD3/CD28 agonistic antibodies for 48 h, before being subjected to transcriptomic profiling. Genes associated with potential early exhaustion, such as *Tox2*, *Ctla2a*, *Prdm1*, and *Batf,* were enriched in DIO T cells (**Fig. 5A**). *Rorc*, a gene induced by signaling of TGF-β and IL-6 (two well-established obesity-associated cytokines(30)) was upregulated in DIO T cells. Next, GSEA was performed to delineate pathways and processes differing between CON and DIO T cells.

**Figure 5.**
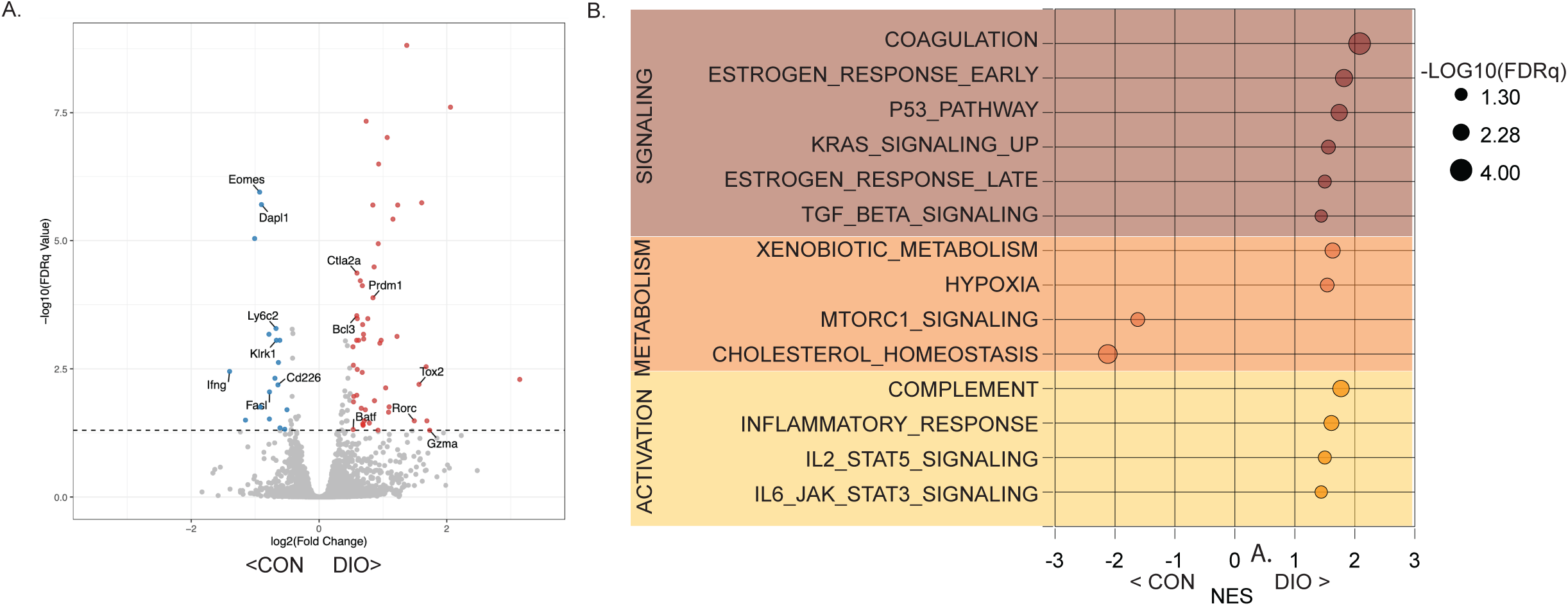
Activated T cells from DIO mice reveal a premature effector signature. Volcano plot of differentially expressed genes (A) and GSEA of transcriptomic profile using Hallmark gene sets (B) in T cells isolated from spleens of CON and DIO mice and incubated for 48 h with anti-CD3/CD28 agonistic antibodies. Normalized enrichment scores (NES) and FDRq values are presented. n=5 mice/group.

MSigDB Hallmark gene sets most enriched in activated T cells from CON mice by GSEA included mTORC1 signaling and cholesterol homeostasis, while gene sets associated with signaling (coagulation, TGF-β, KRAS, etc.) and activation (inflammatory response, IL-2, IL-6 signaling, etc.) were enriched in activated T cells from DIO mice (**Fig. 5B**). Overall, these transcriptomic results suggest that obesity may promote transcriptomic profiles in effector T cells that restrict metabolic adaptation and the maintenance of memory cells.

### The tumor microenvironment in obesity impairs the activity of CAR-T cells

Obesity limits T cell function and promotes an immunosuppressive TME.(19,22) Hence, we first queried whether B7-H3.CAR-T cells generated from age-(25-30 week old) and diet-matched (CON vs DIO) mice would perform similarly in vivo. To do so, we generated B7-H3.CAR-T cells from both CON and DIO mice and infused them into CON and DIO tumor-bearing mice (**Fig. 6A**). We observed more robust tumor control in CON tumor-bearing mice treated with B7-H3.CAR-T cells generated from either CON or DIO mice compared to DIO tumor-bearing mice treated with B7-H3.CAR-T cells generated from either CON or DIO mice (**Fig. 6B, Supp.** Fig. 4A) suggesting that in obese mice CAR-T cells may face a more immunosuppressive TME. Sustained high-fat diet feeding necessitates that DIO studies routinely result in mice of 20-30 weeks of age, during which time age-matched CON mice gain significant weight (**Supp.** Fig. 4B). To determine the extent to which such age-related weight gain might confound our results, we investigated the efficacy of B7-H3.CAR-T cells generated from lean 12-week-old mice when transferred into 25–30-week-old CON or DIO mice (**Fig. 6C**). The adoptive transfer of B7-H3.CAR-T cells obtained from lean mice resulted in robust tumor control in both CON and DIO mice (**Fig. 6D**), leading to no significant differences in survival (**Supp.** Fig. 4C). Together these results indicate that obesity constrains the efficacy of CAR-T cell therapy by both engendering an immunosuppressive TME and by constraining T cell function. However, CAR-T cells generated from lean mice can overcome the inhibitory factors associated with obesity.

**Figure 6.**
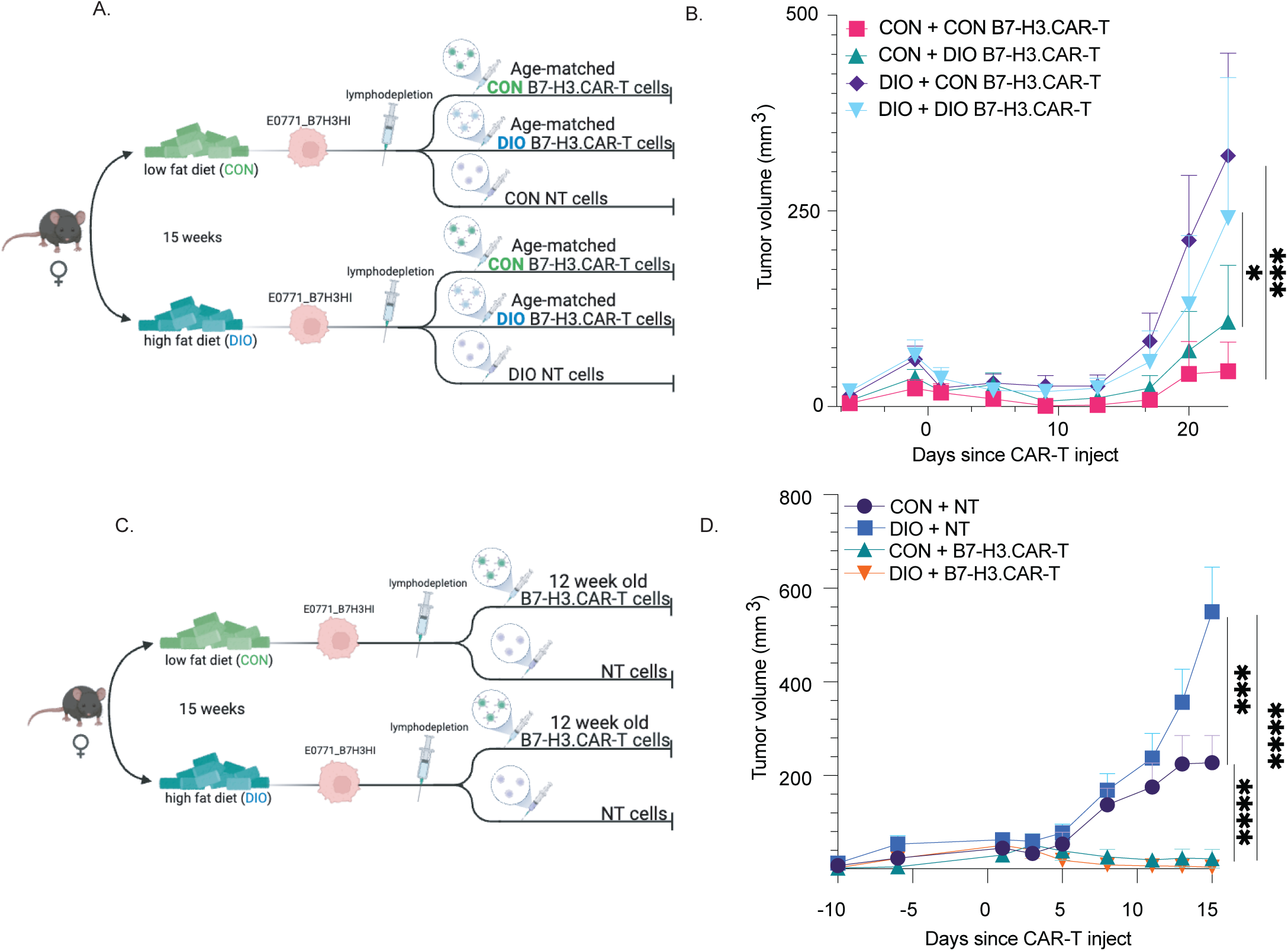
The tumor microenvironment in obesity impairs the activity of CAR-T cells. A. Representative schematic of the tumor model in which female mice placed on low-(CON) or high-fat (DIO) diet for 15 weeks were then engrafted with the E0771_B7H3HI tumor cell line, treated with chemotherapy based lymphodepletion, and infused with age-matched B7-H3.CAR-T cells generated from either CON or DIO mice. B. Measurement of tumor volumes overtime in the tumor model described in (A). C. Representative schematic of the tumor model in which female mice are placed on low-(CON) or high-fat (DIO) diet for 15 weeks before tumor E0771_B7H3HI implantation, treated with chemotherapy based lymphodepletion, and infused with B7-H3.CAR-T cells or NT cells obtained from 12-week-old mice. D. Measurement of tumor volumes overtime in the tumor model described in (C). n = 9-16 mice/group. Significance determined by linear regression. *p < 0.05; ***p < 0.001; ****p < 0.0001.

### Obesity impairs CAR-T cell-induced immunological memory

Clearly CAR-T cells from young lean mice resulted in potent tumor control in both control and obese tumor-bearing mice, but the impact of obesity on immunological memory is unclear. Hence, we next investigated whether obesity could impact CAR-T cell persistence and in particular their ability to become tissue-resident memory (TRM)-like cells, which provide immune surveillance and rapid response to tumor cells.(31,32) CON and DIO tumor-bearing mice treated with B7-H3.CAR-T cells obtained from lean mice (12-week old) that cleared their tumor (defined as no tumor recurrence by day 100) were rechallenged with the same cell line and monitored for an additional 80 days. CON and DIO mice that were naïve to both the E0771_B7-H3HI cells and B7-H3.CAR-T cells served as a control and were also injected with E0771_B7H3HI cells at this time (**Fig. 7A**). Almost all control mice that previously survived CAR-T treatment (CON + B7-H3.CAR-T) exhibited durable protection from tumor rechallenge, with a survival rate of 91%. In contrast, durable protection was evident in only 44% of DIO mice (DIO + B7-H3.CAR-T) (**Fig. 7B**). To further interrogate how obesity impairs durability of CAR-T cell responses, we first isolated and profiled circulating T cells 72 h after tumor rechallenge.

**Figure 7.**
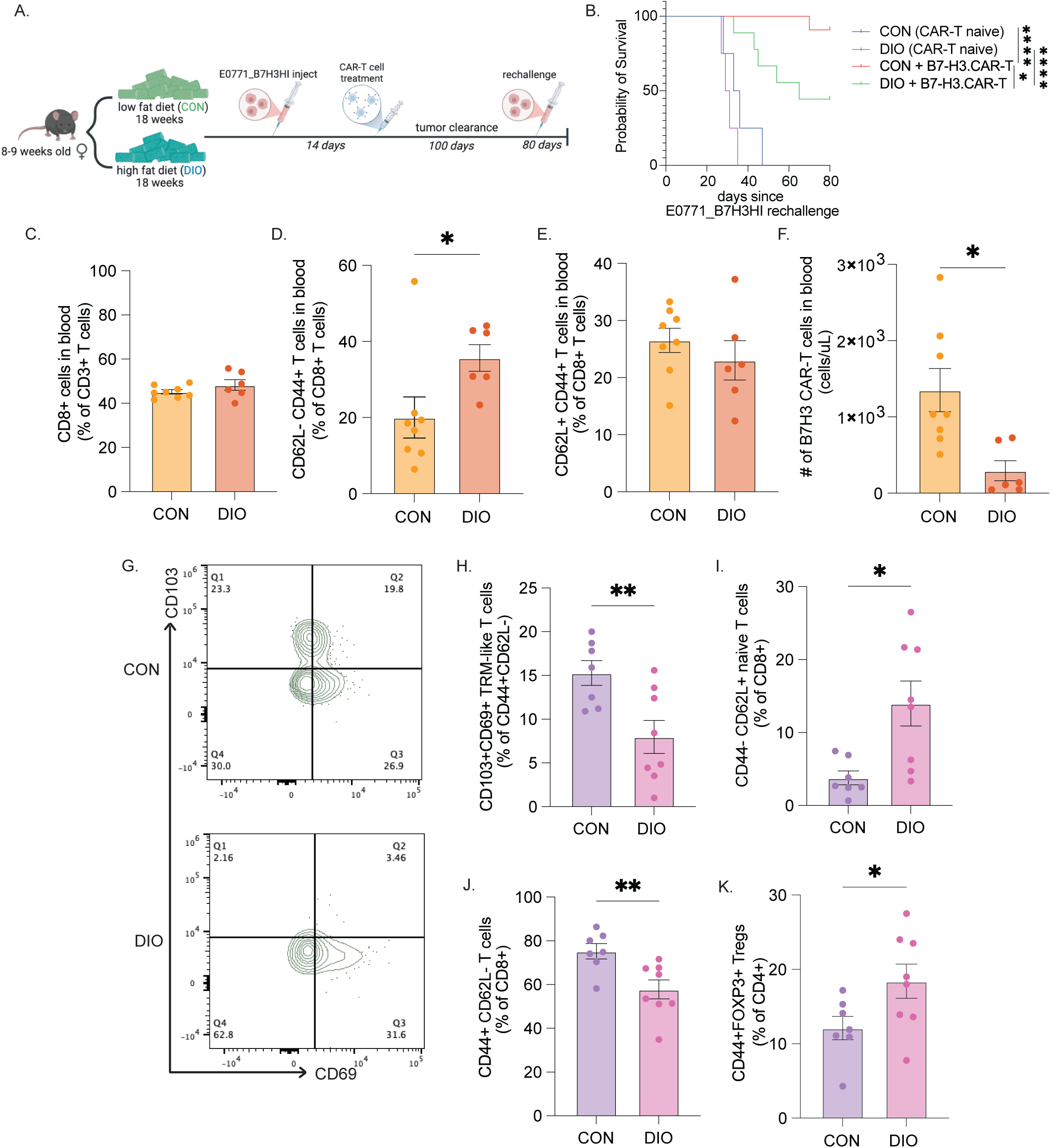
Obesity impairs CAR-T cell-induced immunological memory. A. Representative schematic of the tumor model in which female mice were placed on low-(CON) or high-fat (DIO) diet for 15 weeks before engraftment with the E0771_B7H3HI cell line followed by chemotherapy-based lymphodepletion, and infused of B7-H3.CAR-T cells. Mice that cleared their tumors and remained tumor-free for 100 days were rechallenged with the same cell line and monitored for 80 days for tumor growth. B. Probability of survival after E0771_B7-H3HI rechallenge (n=4-5/group B7-H3.CAR-T naïve control, and 9-11/group B7-H3.CAR-T treated). C-F. CON and DIO mice were bled 72 h after tumor rechallenge to measure the frequency of CD8+ (C), CD8+CD62L-CD44+ (D), CD8+CD62L+CD44+ (E), and B7-H3.CAR-T cells (F) among CON vs DIO PBMCs. G-K. Immune profiling of the fourth mammary fat pad (MFP) of CON and DIO mice that survived initial B7-H3.CAR-T cell treatment. Representative flow cytometry plot (G) and quantification (H) of CD103+CD69+ TRM-like T cells, and quantification of CD44-CD62L+ naive T cells (I), CD44+ CD62L-T cells (J), and CD44+FOXP3+ Tregs (K). n=6-8/group for PBMC data and n=7-8/group for MFP data. Significance determined by Logrank (Mantel-Cox) test (B) and Student’s t-tests (C-K). *p < 0.05; **p < 0.01; ****p < 0.0001.

While no significant difference in total circulating CD8+ T cells was observed (**Fig. 7C**), DIO mice had significantly higher CD8+CD62L-CD44+ circulating cells (**Fig. 7D,E**) and significantly fewer counts of circulating B7-H3.CAR-T cells (**Fig. 7F, Supp.** Fig. 5A) relative to CON mice. We also profiled immune cells isolated from the 4^th^ MFP harvested 100 days after initial CAR-T treatment. No significant differences were observed in CD4+, CD8+ or CD8+CD44+ T cell counts (**Supp.** Fig. 5B-E). In contrast, DIO mice had fewer CD103+ CD69+ TRM-like T cells (**Fig. 7G,H**), indicating that obesity curtailed the establishment of tissue residency. DIO mice had significantly higher CD8+CD44-CD62L+ naïve T cells (**Fig. 7I, Supp.** Fig. 5F) and significantly fewer CD8+CD44+CD62L-T cells (**Fig. 7J**), suggesting obesity may prevent the establishment of the residency immunologic program. Finally, DIO mice showed increased CD44+FOXP3+ regulatory T cells (**Fig. 7K, Supp.** Fig. 5G), which may contribute to the immunosuppressive TME. Together these data indicate that obesity may impair the establishment or maintenance of TRM-like CD8+ T cells and reduce the persistence of CAR-T cells.

## Discussion

Here we demonstrated that B7-H3.CAR-T cell therapy is effective in a syngeneic TNBC model, but obesity blunts the ability of B7-H3.CAR-T cells to elicit durable immunological memory. Furthermore, we identified an obesity-B7-H3 axis that enables accelerated TNBC growth.

We have previously reported an extensive analysis of the expression of B7-H3 in TNBC, which strongly supported the use of B7-H3.CAR-T cells in the attempt to treat TNBC.(7,8) Here, we first aimed to understand if B7-H3 in TNBC may be regulated by cytokines associated with obesity, which may directly impact the capacity of B7-H3.CAR-T cells to recognize and eliminate TNBC cells. Past studies identified that mTORC1 signaling upregulates B7-H3 in a mouse model of renal cancer,(33) and FUT8 fucosyltransferase sustains B7-H3 expression in TNBC.(34) Here we show that, similar to PD-L1,(35) B7-H3 can be upregulated via inflammatory signaling of cytokines-associated with obesity in TNBC cells. Importantly, unlike previous reports demonstrating that suppression of B7-H3 limits tumor growth in control mice,(33) we show that B7-H3 expression is pro-tumorigenic in DIO mice. This association between B7-H3 expression, obesity, and tumor growth is particularly relevant because it highlights that targeting B7-H3 may prevent tumor escape due to its critical role in tumor growth in the specific proinflammatory environment associated with obesity. However, it remains to be demonstrated if the obesity-B7-H3 axis observed here is restricted to the cytokine milieu of obesity or if other proinflammatory conditions might result in similar acceleration of tumor growth by B7-H3.

Epidemiological studies suggest that obesity has little impact on the clinical outcome of CAR-T therapy for blood cancers.(36,37) However, obesity is well established to attenuate the formation of memory-like T cells in influenza(38,39) and other vaccinations (e.g., COVID-19(40)), despite similar initial responses to vaccination. Thus, it is critical to explore the role of obesity in adoptive immune therapy models to better understand and identify possible mechanistic links.

Using a well-controlled preclinical model, we report several critical observations on how obesity can impact the functionality of CAR-T cells. The first observation is that obesity intrinsically affects the molecular adaptation of T cells to activation. Specifically, circulating T cells obtained from DIO mice and activated ex vivo demonstrated significant upregulation of genes associated with effector function (*Batf, Prdm1,* and *Tox2*) compared to T cells obtained from CON mice. Although how obesity imprints circulating T cells remains to be explored mechanistically, it is important to note the dysfunction of T cells from DIO mice. Specifically, CAR-T cells from DIO mice retain similar effector function but harbor impaired metabolic fitness and skewed induction of memory features after encountering tumor cells in vitro compared to CAR-T generated from CON mice. The second observation is that obesity impairs the antitumor effects of CAR-T cells even if DIO mice are treated with CAR-T cells generated from age-matched CON mice indicating that potent CAR-T cells generated ex vivo can still be inhibited by factors associated with obesity in vivo. Finally, we observed that CAR-T cells generated from young lean mice can adequately control the tumor growth in DIO mice but fail in controlling tumor rechallenge, indicating that obesity has a prominent effect in preventing memory formation.

TRM-like T cells are critical for long term protection in tumor models.(32) In our model we observed that obesity profoundly impairs TRM-like T cell development, a key aspect of memory formation. To become TRM-like, T cells must undergo a series of transcriptional, phenotypic, and functional shifts.(32) The critical role played by TRM-like T cells in cancer has emerged very recently; thus there is scant evidence as to how obesity might mechanistically alter TRM-like T cell differentiation which will require additional studies. Furthermore, our study is limited to a specific tumor model, thus it remains to be demonstrated if the observation can be recapitulated in other solid tumor models and other CAR-T cell targets. We generated a cell line that constitutively overexpressed human B7-H3. While this model was essential to answer our current research question, genetic knock-in of human B7-H3 to the mouse allele would allow for future interrogation of interactions between CAR-T cell therapy and native B7-H3 expression.

In summary, we have shown that obesity potently suppresses multiple facets of CAR-T cell efficacy, and that the newly identified association of obesity and B7-H3 expression holds significant potential as a target for interventions to limit obesity-driven cancer. Moreover, our data demonstrate that a failure of adequate TRM-like T cell differentiation contributes to impaired durability of responses achieved with CAR-T cell therapy in obesity. Additional studies, including adoptive transfer of TRM-like T cells from control mice, will be necessary to confirm this mechanism. Taken together, we show that obesity is an important but understudied determinant of CAR-T cell therapy response.

## Acknowledgements

We would like to acknowledge Dr. Jessica Thaxton, Dr. Justin Milner, and Dr. William Green for their thoughtful feedback and support. Some summary figures were created with Biorender.com. This work was enabled by the Lineberger Preclinical Research Unit and the Flow Cytometry Core Facility at the University of North Carolina at Chapel Hill, which are supported in part by an NCI Center Core Support Grant (P30 CA16086) to the UNC Lineberger Comprehensive Cancer Center. Funding for the project was provided by a grant from the Breast Cancer Research Foundation (to SDH) and the UNC Triple Negative Breast Cancer Center (to SDH and GD).

## Acknowledgements

*This work was enabled by the Lineberger Preclinical Research Unit and the Flow Cytometry Core Facility at the University of North Carolina at Chapel Hill, which are supported in part by an NCI Center Core Support Grant (P30 CA16086) to the UNC Lineberger Comprehensive Cancer Center*.

