## Supplemental figures for "Obesity Impairs the Antitumor Activity of CAR-T Cells in Triple-Negative Breast Cancer"

Supplementary figure 1

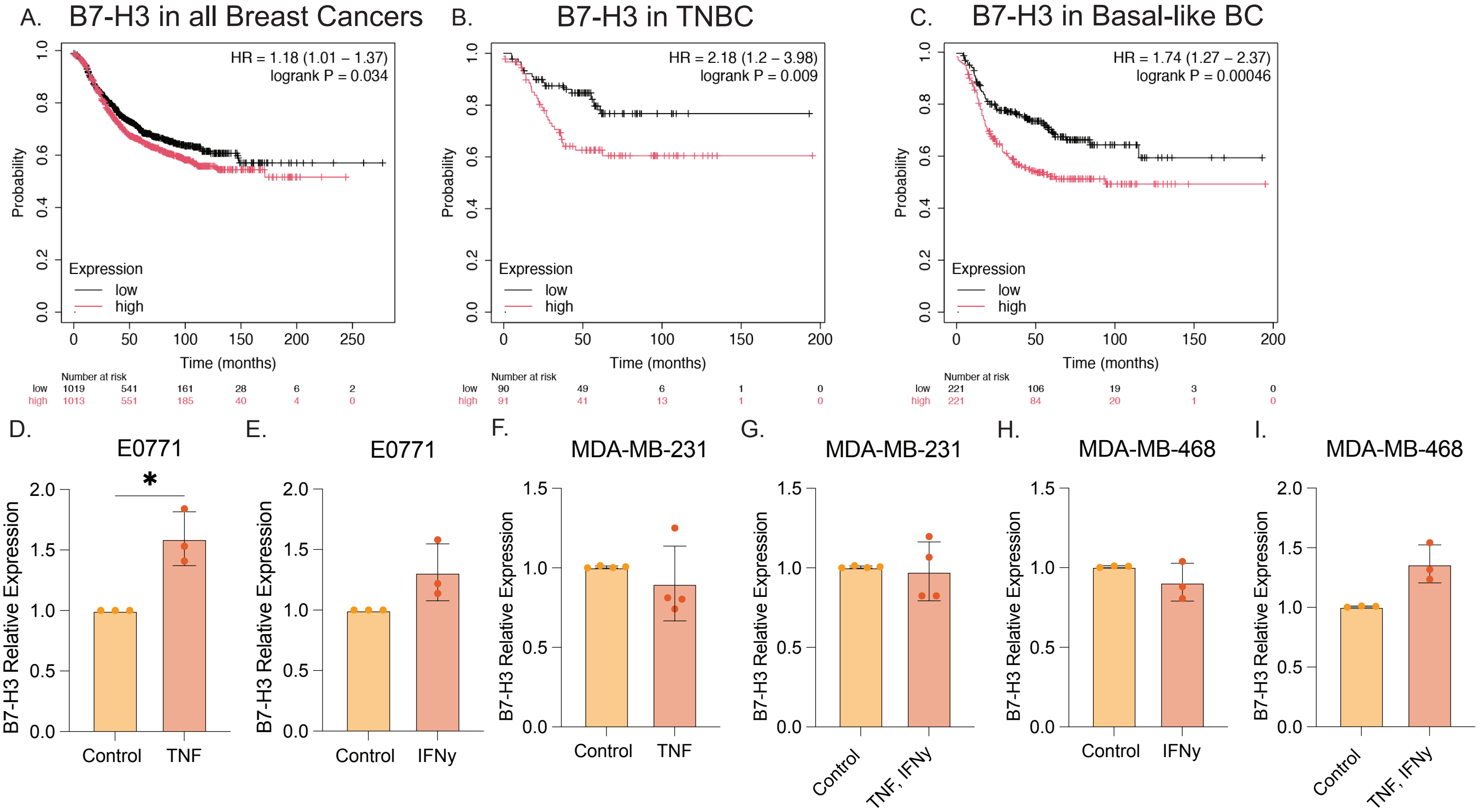


Kaplan-Meier plots of all breast cancers (A), triple-negative breast cancers (B) and basal-like breast cancers (C) sorted by high or low CD276 expression retrieved from KMPlotter.com. qPCR quantification of B7-H3 in E0771 cells treated with TNF (D), IFNγ (E), or control. qPCR quantification of B7-H3 in MDA-MB-231 (F-G) or MDA-MB-468 (H-I) cells treated with TNF alone, in combination with IFNγ, or control. n=3-4 group. Significance determined by Student's t-test. $*p < 0.05$; $**p < 0.01$.

Supplementary figure 2

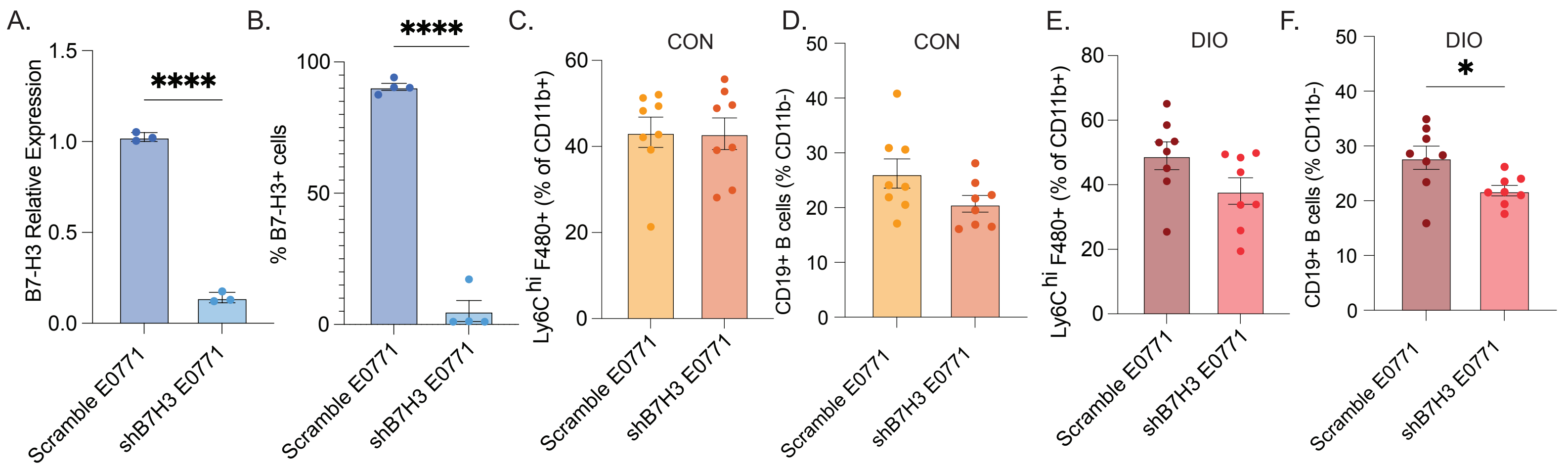


A. qPCR results measuring relative expression of B7-H3 in scramble E0771 and shB7H3 E0771 cells. B. Flow cytometry measurement of surface expression of B7-H3 in scramble E0771 and shB7H3 E0771 cells. C. Ly6ChiF480+ macrophages (%CD11b+) from tumors in CON mice D. CD19+ B cells (%CD11b-) from tumors in CON mice. E. Ly6ChiF480+ macrophages (%CD11b+) from tumors in DIO mice. F. CD19+ B cells (%CD11b-) from tumors in DIO mice. n=3/4 per group (A,C) 7/8 per group (C,D). Significance determined by Student's t-test (A-F),(*p < 0.05; ****p < 0.0001).

Supplementary figure 3

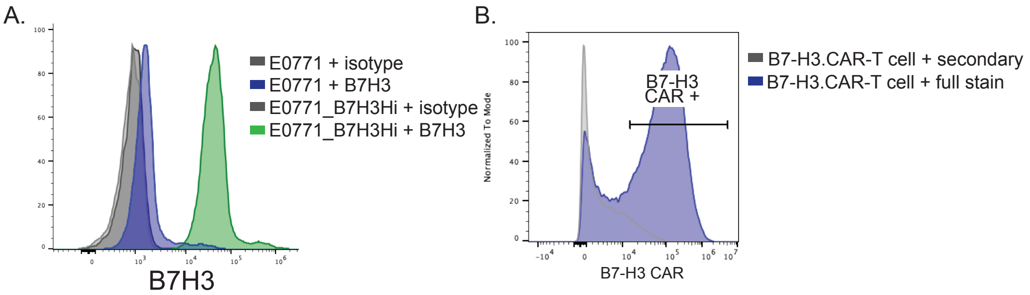


A. Representative flow plot of transduction and sorting of E0771_B7H3HI. B. Representative flow plot of transduction of T cells with B7-H3.CAR.

Supplementary figure 4

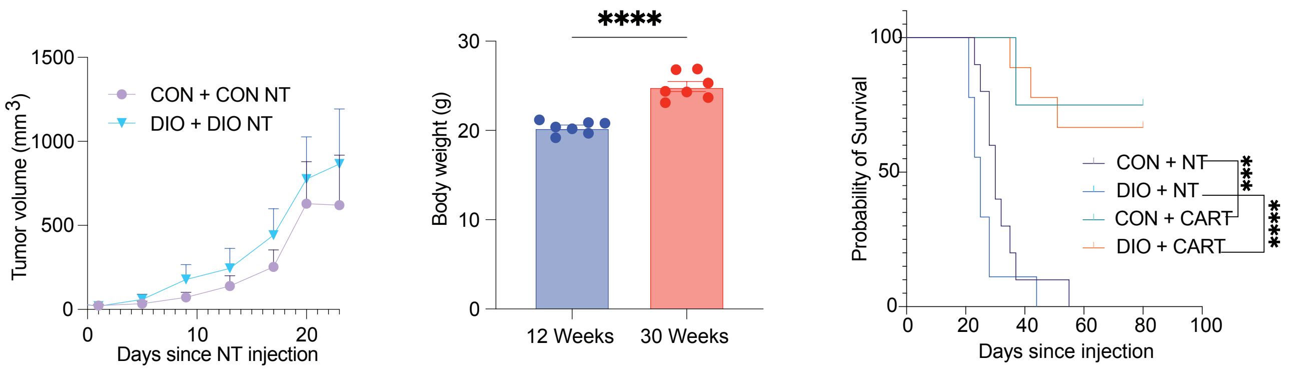


A. Tumor volume over time for CON or DIO mice receiving NT cells monitored with calipers over 23 days (n=8/group). B. Body weights of mice on control diet (CON) at 12 or 30 weeks of age. n= 8 mice/group. C. Survival plot of CON or DIO mice receiving 12-week-old CAR-T cells monitored over 15 days. n = 9-16 mice/group. Significance determined by linear regression (A) Student's t-test (B), Logrank (Mantel-Cox) test (C) (***p < 0.001; ****p < 0.0001).

Supplementary figure 5

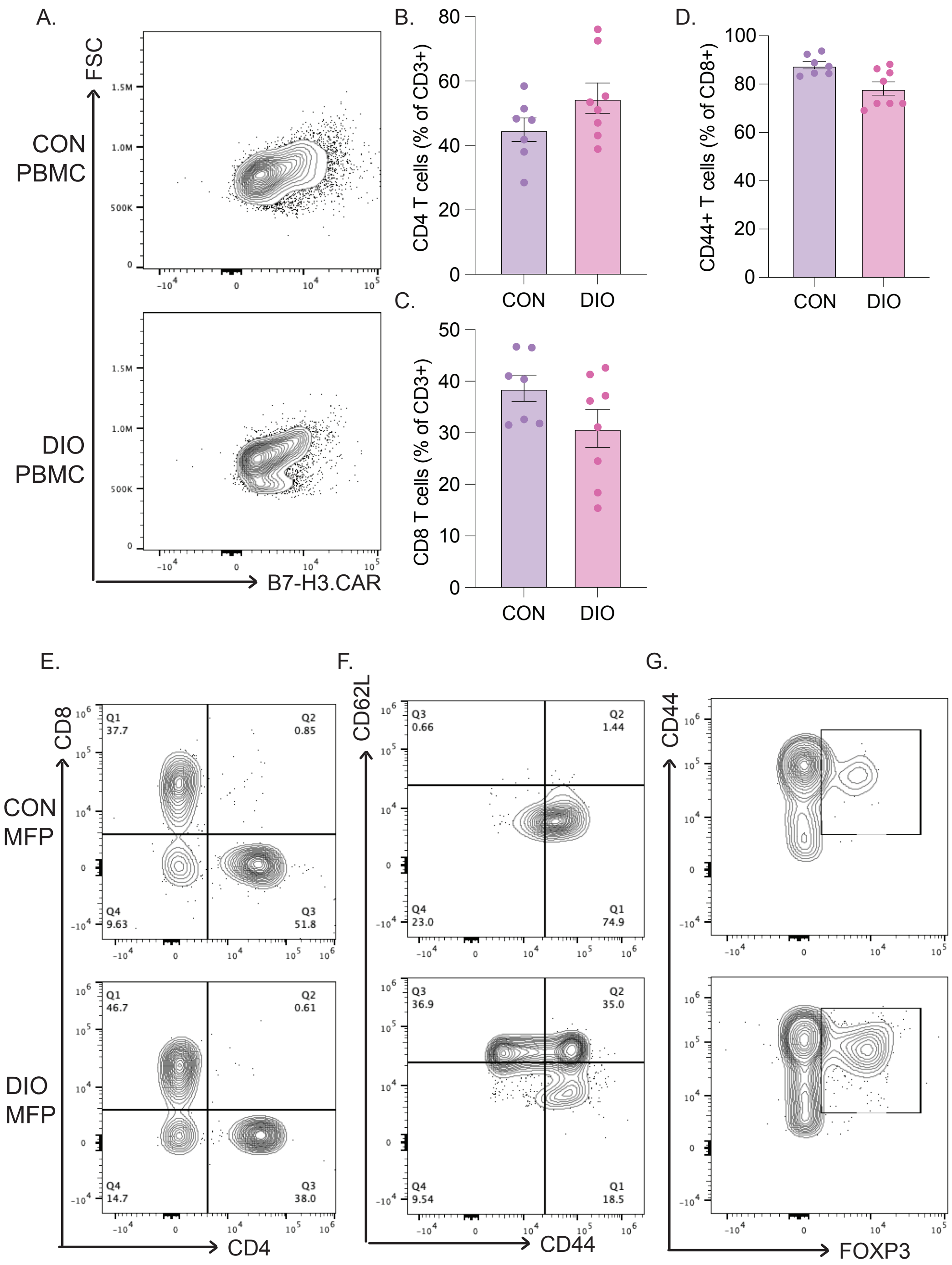


A. Representative flow plot for PBMCs isolated from CON or DIO mice, stained for B7-H3.CAR. B-D. CD4+ T cells (% CD3+) (B), CD8+ T cells (%of CD3) (C), CD44+ T cells (% CD8+) (D) from MFP of CON and DIO mice. E. Representative flow plot for CD8/CD4 T cells isolated from MFP of CON or DIO mice. F-G. Representative flow plots for CD62L/CD44 T cells (F) and CD44/FOXP3 T cells (G) isolated from MFP of CON or DIO mice. Significance determined by student’s t-test (B-D).
