## Supplemental table for "Obesity Impairs the Antitumor Activity of CAR-T Cells in Triple-Negative Breast Cancer"

Supplementary table 1: Antibodies used for flow cytometry.

| **Marker** | **Dilution** | **Supplier** | **Cat #** |
|---|---|---|---|
| BCL2 | 600 | BioLegend | 633507 |
| CD103 | 500 | BioLegend | 121433 |
| CD11b | 200 | eBioscience | 46-0112-82 |
| CD11c | 200 | BD | 564986 |
| CD19 | 150 | BioLegend | 115531 |
| CD206 | 200 | BioLegend | 141721 |
| CD3 | 200 | BioLegend | 100216 |
| CD4 | 200 | BD | 741913 |
| CD44 | 400 | BD | 741227 |
| CD45 | 300 | BD | 564279 |
| CD62L | 200 | BioLegend | 104433 |
| CD69 | 400 | BioLegend | 109-606-088 |
| CD8a | 200 | BioLegend | 100758 |
| F4/80 | 200 | BioLegend | 123124 |
| FoxP3 | 200 | eBioscience | 17-5773-80A |
| GzmB | 200 | BioLegend | 372225 |
| LAG3 | 200 | BioLegend | 125226 |
| Ly6C | 300 | BioLegend | 128055 |
| Ly6G | 150 | BioLegend | 127633 |
| MHCII | 200 | BioLegend | 107615 |
| PD-1 | 200 | BioLegend | 135224 |
| TIM3 | 200 | BD | 747618 |
| TOX | 100 | Miltenyi | 130-120-716 |
| Viability | 1500 | Invitrogen | L34961 |
